# An autism-associated calcium channel variant causes defects in neuronal polarity and axon termination in the ALM neuron of *C. elegans*

**DOI:** 10.1101/2020.09.11.292003

**Authors:** Tyler Buddell, Christopher C. Quinn

## Abstract

Variants of the *CACNA1C voltage-gated calcium channel* gene have been associated with autism and other neurodevelopmental disorders including bipolar disorder, schizophrenia, and ADHD. The Timothy syndrome mutation is a rare *de novo* gain-of-function variant in *CACNA1C* that causes autism with high penetrance, providing a powerful avenue into investigating the role of *CACNA1C* variants in neurodevelopmental disorders. In our previous work, we demonstrated that an *egl-19(gof)* mutation, that is equivalent to the Timothy syndrome mutation in the human homolog *CACNA1C,* can disrupt termination of the PLM axon in *C. elegans*. Here, we find that the *egl-19(gof)* mutation disrupts the polarity of process outgrowth in the ALM neuron of *C. elegans*. We also find that the *egl-19(gof)* mutation can disrupt termination of the ALM axon. These results suggest that the Timothy syndrome mutation can disrupt multiple steps of axon development. Further work exploring the molecular mechanisms that underlie these perturbations in neuronal polarity and axon termination will give us better understanding to how variants in *CACNA1C* contribute to the axonal defects that underlie autism.

## Description

The *egl-19* gene in *C. elegans* encodes the pore forming subunit for the L-type voltage gated calcium channel that is homologous to the *CACNA1C* gene in humans (Lee et al., 1997). Variants in *CACNA1C* are risk factors for autism and other neurodevelopmental disorders (Li et al., 2015; Lu et al., 2012; Strom, et al., 2010). Timothy syndrome is a syndromic form of autism that can be caused by either of three rare *de novo* mutations in *CACNA1C*. These mutations cause either a G402R, G402S or G406R mutation in the CACNA1C protein (Splawski et al., 2004; Bader et al., 2011).

Our previous work demonstrated that PLM axon termination is disrupted by mutations equivalent to the G402R and G406R mutations in CACNA1C (Buddell et al., 2019). Our study also revealed behavioral defects in these mutant worms. Although the anatomical basis for these behavioral defects has not been determined, it is likely that they are caused by multiple defects within the mechanosensory system. Here, we examine other neurons in the mechanosensory system of *egl-19(gof)* mutants and identify defects in the polarity of the ALM neuron as well as defects in the termination of its axon.

The mechanosensory neurons in *C. elegans* are responsible for transducing light touch and consist of two ALM neurons, two PLM neurons, one AVM neuron and one PVM neuron (Chalfie et al., 1985). To identify neuronal defects caused by the *egl-19(gof)* mutation, we labeled each of the six mechanosensory neurons with a fluorescent transgene that is expressed in each of the six mechanosensory neurons. In addition to the previously reported axon termination defects in the PLM neuron, we observed two novel defects in the ALM neuron. In wild-type animals, the cell bodies of the ALM neurons reside on the lateral body wall and extend a single axon into the head, where they terminate prior to reaching the tip of the nose (Figure 1A). In *egl-19(gof)* mutants, we observed overextended ALM axons, where the axon extended past its normal termination point and terminated within the tip of the nose (Figure 1 B,C). We also observed defects in the polarity of the ALM neuron. In wild-type animals, nearly all ALM neurons extend a single process from the cell body (Figure 1D,F). However, in *egl-19(gof)* animals, we often observed a second short process that extended in the posterior direction (Figure 1E,F).

**Figure 1.**
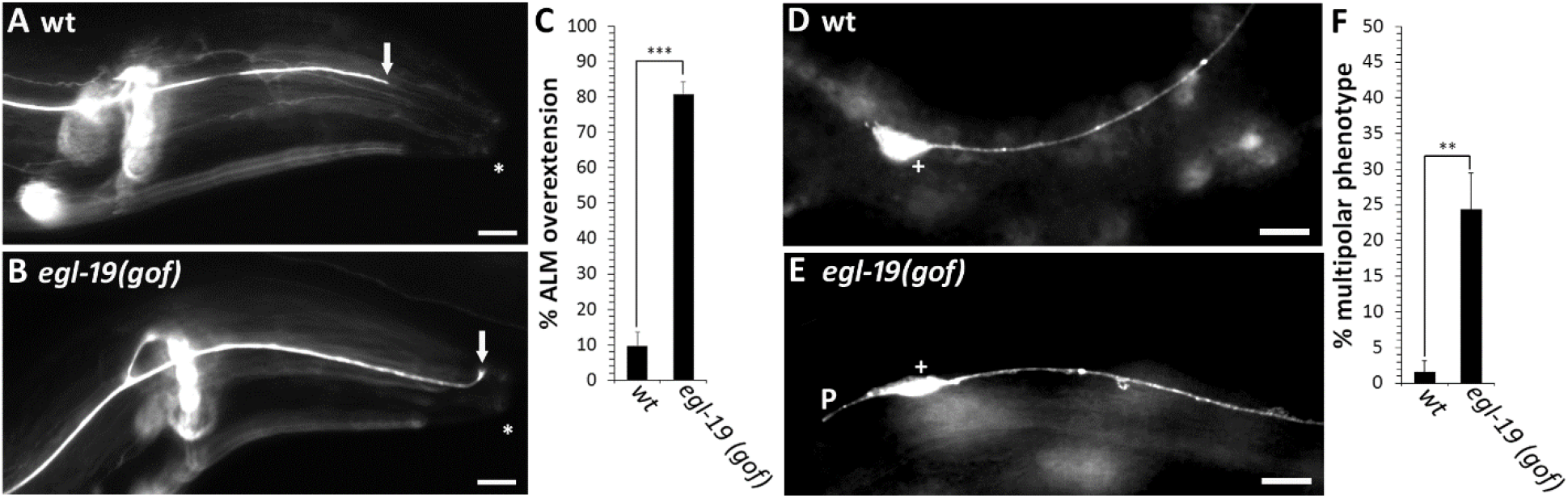
The *egl-19(gof)* Timothy syndrome mutation causes defects in axon termination and neuronal polarity. (A)Example of normal axon termination in a wild-type ALM neuron, where the axon terminates posterior to the mouth of the worm (arrow). (B) Example of axon termination defect in a *egl-19(gof)* mutant, where the ALM axon extends to the anterior most point of the worm (arrow). (C) Quantification of axon overextension defects in ALM neurons. (D) Example of a normal cell body of a wild-type ALM neuron, where a single process extends from the anterior side of the ALM cell body. (E) Example of a multipolar phenotype in a *egl-19(gof)* mutant, where a short process extends from the posterior side of the ALM cell body. (F) Quantification of the multipolar phenotype that is caused by the egl-19(gof) mutation. Axons are visualized with the *muIs32* transgene that encodes *Pmec-7::gfp*. Arrows point to the tip of the ALM axon. Asterisk marks the anterior-most part of the worm. Cross indicates ALM cell body. P indicates a multipolar defect. Scales bars are 10um. Between 100 and 150 axons were observed in L4 stage hermaphrodites per genotype. Asterisks indicate statistically significant difference, Z-test for proportions (**p<0.0005; ***p<0.0001). Error bars represent the standard error of the proportion.

These results suggest that the Timothy syndrome mutation can disrupt multiple steps of axon development. First, the *egl-19(gof)* mutation can disrupt the polarization of process outgrowth. Second, the *egl-19(gof)* mutation can also disrupt ALM axon termination. Future work will address the molecular mechanisms that underlie these alterations in neuronal polarity and axon termination. An understanding of these mechanisms will be critical to our understanding of how variants in *CACNA1C* give rise to the axonal defects that underlie autism.

## Methods

*C. elegans* strains were cultured and maintained on nematode growth medium (NGM)-agar plates using standard methods at 20°C (Brenner, 1974). The following alleles were used in this study: wild-type *N2; egl-19(n2368)*. These strains were obtained from the CGC. For analysis of axon termination phenotypes, animals were mounted on a 5% agarose pad and observed with a 40x objective. For PLM axon termination, an axon was scored as defective if it grew anterior to the ALM cell body. PLM & ALM neurons were visualized with the *muIs32* transgene which encodes Pmec-7::gfp + lin-15(+) and is expressed in all mechanosensory neurons (Ch’ng et al., 2003). The *muIs32* transgene was obtained from the CGC. The microscope used for imaging and phenotype analysis was the Zeiss Axio Imager M2. Images were acquired using the AxioCam MRm camera. Fluorescence was illuminated using the X-cite series 120Q. Images were taken under a 40x objective unless otherwise specified. All images acquired from the microscope were analyzed using Axiovision 4 software. Images were edited into figures using Adobe Photoshop.

## Acknowledgements

We would like to thank the Caenorhabditis Genetics Center for strains.

## Author Contributions

Tyler Buddell: Investigation, Data curation, Formal Analysis, Writing - original draft, Writing - review and editing. Christopher C. Quinn: Investigation, Funding acquisition, Supervision, Writing - review and editing.

